# Emergence of an adaptive command for orienting behavior in premotor brainstem neurons of barn owls

**DOI:** 10.1101/119180

**Authors:** Fanny Cazettes, Brian J. Fischer, Michael V. Beckert, Jose L. Pena

## Abstract

The midbrain map of auditory space commands sound-orienting responses in barn owls. Owls precisely localize sounds in frontal space but underestimate the direction of peripheral sound sources. This bias for central locations was proposed to be adaptive to the decreased reliability in the periphery of sensory cues used for sound localization by the owl. Understanding the neural pathway supporting this biased behavior provides a means to address how adaptive motor commands are implemented by neurons. Here we find that the sensory input for sound direction is weighted by its reliability in premotor neurons of the owl’s midbrain tegmentum such that the mean population firing rate approximates the head-orienting behavior. We provide evidence that this coding may emerge through convergence of upstream projections from the midbrain map of auditory space. We further show that manipulating the sensory input yields changes predicted by the convergent network in both premotor neural responses and behavior. This work demonstrates how a topographic sensory representation can be linearly read out to adjust behavioral responses by the reliability of the sensory input.

**Significance statement:** This research shows how statistics of the sensory input can be integrated into a behavioral command by readout of a sensory representation. The firing rate of midbrain premotor neurons receiving sensory information from a topographic representation of auditory space is weighted by the reliability of sensory cues. We show that these premotor responses are consistent with a weighted convergence from the topographic sensory representation. This convergence was also tested behaviorally, where manipulation of stimulus properties led to bidirectional changes in sound localization errors. Thus a topographic representation of auditory space is translated into a premotor command for sound localization that is modulated by sensory reliability.

## INTRODUCTION

The synthesis of motor commands weighted by sensory reliability is critical for understanding adaptive behavior. To address this question, we investigated the neural basis of the centrality bias of barn owls, which is a systematic underestimation of sound sources towards the center of gaze (Hausmann et al., 2009; Knudsen et al., 1979). This centrality bias may be optimal for the owl’s prey capture, where the amount of bias reflects the relative weight given to sensory evidence depending on the reliability of the sensory input (Fischer and Pena, 2011; Girshick et al., 2011; Schwartz et al., 2007). Yet, the neural circuit decoding a behavioral command that integrates the reliability of sensory evidence has not been demonstrated. This study describes how such a command may emerge in a premotor neural population that controls orienting behaviors in the sound localization system of the owl.

Most animals use interaural time difference (ITD) for inferring the horizontal direction of sound location (Strutt, 1907; Blauert, 1997). In the barn owl, a map of auditory space emerges in the external nucleus of the inferior colliculus (ICx) (Knudsen and Konishi, 1978), where selectivity for ITD determines the horizontal spatial tuning of auditory neurons. The ICx map is a distorted representation of space where the front is overrepresented (Knudsen, 1982)(Figure 1). Additionally, frequency and spatial selectivity of ICx neurons co-vary (Cazettes et al., 2014), where neurons preferring frontal directions are tuned to higher frequencies than peripheral neurons (Figure 1). This direction-dependent frequency selectivity corresponds to how the variability of interaural phase difference changes across frequency and direction, estimated from the owl’s head-related transfer function (Cazettes et al., 2014). Previous studies have proposed that this non-uniform spatial and frequency selectivity across the ICx allows for the representation of ITD reliability (Fischer and Pena, 2011; Rich et al., 2015; Cazettes et al., 2016).

**Figure 1.**
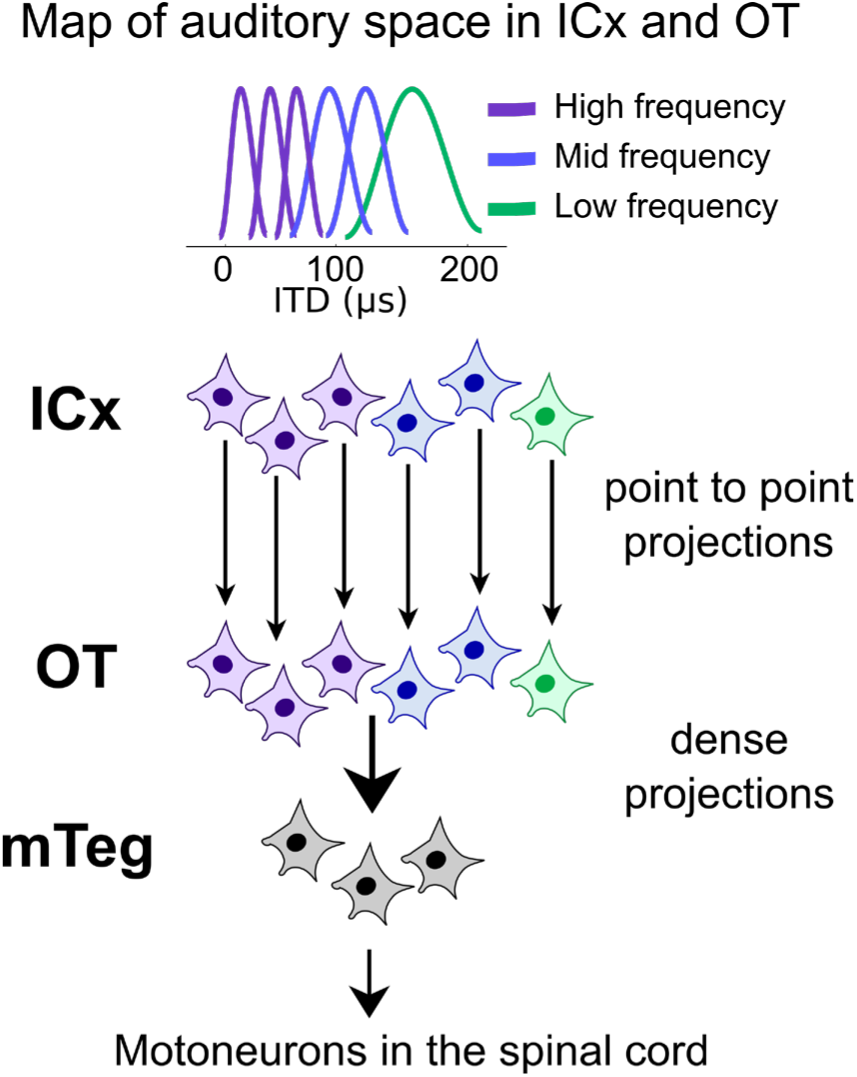
Sound localization pathway. ICx and OT display maps of frequency and space (Knudsen, 1982; Cazettes et al. 2014). Frontal space (small ITD in μs in purple) is associated with high frequency. Conversely, peripheral space (large ITD in μs in green) is associated with low frequency. Although OT directly projects to mTeg, the nature of the projection between OT and mTeg is unknown.

ICx projects point-to-point to the optic tectum (OT), analog of mammalian superior colliculus (Knudsen and Knudsen, 1983), leading to both structures having neurons with similar spatial and frequency tunings (Olsen et al., 1989; Knudsen, 1984). As a result, both ICx and OT display similar maps of auditory space, here referred to as the space-map (Figure 1). OT projects to the midbrain tegmentum (mTeg), which drives the spinal-cord motoneurons that generate the head-orienting response to sound (Masino and Knudsen, 1992, 1993) (Figure 1). In vertebrates, the tectotegmental pathway contributes prominently to the generation of orienting movements of the eyes, head, and body (Masino, 1992; du Lac and Knudsen, 1990). Anatomical studies of the tectotegmental projections in the barn owl suggest that the topographic representation of space in the OT is translated into a representation of orthogonal movement vectors in mTeg (Masino and Knudsen, 1990, 1992, 1993).

This study examined whether responses of premotor neurons in mTeg match the owl’s sound localizing behavior, how these responses emerge, and whether they reflect the reliability of the sensory input. We found the mean population response in mTeg matches the owl’s behavior, and could be explained by a convergence from OT to mTeg. Furthermore, both the mTeg responses and orienting bias of behaving owls changed with manipulations of the sensory reliability and sound frequency in a consistent manner. Thus, we provided evidence of a neural network that weights sensory input by its reliability to encode an adaptive behavioral command.

## MATERIALS & METHODS

### Electrophysiology

#### Surgical procedure

Surgery was performed as described previously (Wang et al., 2012). Briefly, adult American barn owls (*Tyto furcata*; 3 females and 3 males; ages 3 to 5 years old) were anesthetized with intramuscular injections of ketamine hydrochloride (Ketaset; 20 mg/kg, intramuscular) and xylazine (Anased; 4 mg/kg, intramuscular). Prophylactic antibiotics (ampicillin; 20 mg/kg, intramuscular) and lactated Ringer’s solution (10 ml, subcutaneous) were injected at the beginning of each session. An analgesic, carprofen (Rimadyl; 3mg/kg, intramuscular), was administered at the end of each session to prevent inflammation and pain. These procedures complied with National Institutes of Health and guidelines of the Albert Einstein College of Medicine’s Institute of Animal Studies.

Recording from awake animals was impractical in these experiments as electrophysiology required serial electrode penetrations, head restraint, and long recording sessions (up to 10 hours). To control for varying levels of anesthesia, records of the time of each recording relative to anesthetic administration were rigorously maintained. Experiments were interrupted if first-spike latencies or response thresholds changed. Regardless, the effect of anesthesia in the midbrain has been shown to be less pronounced than in the forebrain (Ter-Mikaelian et al., 2007; Schumacher et al., 2011) and therefore lessens concerns about changes in neural response under anesthesia.

#### Extracellular recordings of ICx, OT and mTeg neurons

Single-unit responses were recorded using 1MΩ tungsten electrodes (A-M Systems; Carlsborg, WA). Tucker Davis Technologies (TDT; Alachua, FL) System 3 and custom written Matlab software (Mathworks) were used to record neural data. ICx and OT were located stereotaxically and confirmed by the characteristic responses to ITD and interaural level difference and firing properties (Wang et al., 2012). The mTeg was also located stereotaxically (Masino and Knudsen, 1992, 1993) but recording from this structure was further verified with microstimulation (see below).

#### Microstimulation

Recordings in mTeg were confirmed with microstimulation. At the end of each recording, microstimulation at the recording site was conducted to verify that eye movements were elicited and that the same electrical stimulus was no longer able to elicit eye movements when the electrode was moved a few hundred micrometers away from the recording site. A programmable digital stimulator (WPI, DS8000) was used to deliver trains of current pulses (5ms pulse width, 200Hz, 80ms duration) through the recording electrode, varying the current amplitude from 50 to 150 μA with inter-stimulus intervals larger than 15 seconds (as described in Netser et al., 2010). In most experiments, 130 μA or less was enough to induce unequivocal eye movements.

### Sound stimulation

All experiments were performed inside a double-walled sound-attenuating chamber. Custom-made earphones consisting of a calibrated speaker (Knowles model 1914) and a microphone (Knowles model EK-23024) were inserted in the owl’s ear canal (Wang et al., 2012). Auditory stimuli delivered through the earphones consisted of 100 to 150 ms signals with a 5 ms rise-fall time at 10-20 dB above threshold. For each ICx, OT, and mTeg neuron, we first measured the ITD and rate-level responses with broadband noise (0.5 – 10 kHz). ITD was initially varied from ±300 μs with 30 μs steps over 5-10 trials for ICx and OT and 20 trials for mTeg. Sound level varied from 0 dB SPL to 70 dB SPL in 5-dB steps. Frequency tuning was measured with tones ranging from 500 Hz to 8000 Hz varied in 200 to 500 Hz steps over 20 trials. Neural responses to ITD in mTeg were also recorded at various frequencies (from 1 to 6 kHz, 1 kHz steps, same ITD protocol as broadband). Stimuli within all tested ranges were randomized during data collection. To manipulate the binaural correlation (BC), three random noise signals (N1, N2, and N3) were generated. N1 was delivered to one ear and its copy with a time shift, to produce an ITD, was delivered to the other ear. N2 and N3 were added to the N1 in the left and right ear respectively. These additional noise signals reduced the correlation between the signals in the two ears, depending on the relative amplitudes of the uncorrelated and correlated noises. Binaural correlation was calculated from BC = 1/(1 + k^2^), where k is the ratio between the root-mean-square amplitudes of the uncorrelated and correlated noises (Saberi et al., 1998). On a subset of ICx neurons (n=46) and on all recorded mTeg neurons, we obtained ITD curves (±300 μs; 30 μs steps; 20 repetitions) using noise bursts at different BCs (from 0 [noise in each ear is independent] to 1 [noise in each ear is the same], in 0.2 steps).

Finally, for all mTeg neurons we measured ITD curves (±300 μs; 30 μs steps; 20 repetitions) with low (500-3500 Hz) and high (4500-7500 Hz) frequency band-pass sounds as well as broadband noise (500-9000 Hz).

### Behavior

#### Remote eye tracking

Eye movements triggered by microstimulation in mTeg were measured with high-speed video imaging using a commercially available eye tracking system (EyeLink 1000) that detected the pupil and the position of the corneal reflection. Eyelids were opened momentarily to conduct the test after each recording using a custom made retractor. This procedure and the anesthesia did not prevent blinking of the nictitating membrane, allowing for natural protection of the eyes from drying.

#### Head-orienting behavior

Head-orienting behavior to sound was measured for three hand-raised barn owls only trained to stand on a perch at the center of a speaker array. The speaker array consisted of an azimuthal linear array of 21 speakers, a subset of a hemispherical array of speakers constructed inside a sound-attenuating chamber (Wang et al., 2012; Cazettes et al., 2014). The linear array spanned 100 degrees to the left and right of the midline, with 10 degrees separation between speakers. Each speaker was calibrated in place using a microphone mounted on a pan-tilt robot located at the array center. Head saccades were recorded with a pair of high-speed infrared cameras (Fastec Imaging, 100 frames per sec) mounted in the corners of the room. Head saccades were then tracked using custom designed image-analysis code (Xcitec, Proanalyst) from images calibrated in 3-D using a grid located at the same position of the owl’s head. Owls were presented with the same noise bursts used to measure neural responses (100 ms long, 5 ms rise-fall time, high and low frequency band-pass sounds, and broadband-noise) but through this free-field array. The setup allowed us to measure head saccades from any starting head position, which made it possible to rule out positional biases. To account for the position of the animal in the field of view of the cameras, the owl’s perch was designed such that it only allowed the bird a single stance. However, owls did not always look straight ahead when the stimulus was presented; therefore, the measured angles of head turn were transformed into a head-centered coordinate system. Owls were unrewarded in this task to avoid biases introduced by training. For each sound presentation, the frequency band (0.5-3.5 kHz, 4.5-7.5 kHz or 0.5-9 kHz) and location (±100 deg, 10 deg separation) of the source were randomized by the software. The experimenter initiated a trial when the owl was standing upright and its beak (the tracking reference) was in the field of view of both cameras. The experimenter remained blind to the properties of the sound.

### Experimental Design and Statistical Analysis

No significant differences between sexes were observed in neural responses; therefore data from the two groups were pooled.

#### Electrophysiological data

For each stimulus parameter, a rate curve was computed by averaging the firing rate during sound stimulation over trials. The spontaneous firing rate, obtained when no sound stimulation was presented, was subtracted from the response to the stimulus. Best ITDs were defined as the center of the range that elicited 50% of the maximum response of each tuning-curve. To assess the relative broadness of the frequency tuning we calculated the areas under the normalized frequency tuning curves. To find the peak of ITD-tuning curves of mTeg neurons to tonal stimulation we first fitted the curves with a cosine function. We then defined the ITD corresponding to the closest maximum from 0 μs as the peak (Cazettes et al., 2014). The changes of ITD tuning with BC were consistent in both hemispheres (maximal changes for sound directions within the contralateral hemifields). Thus, analyses of the mean mTeg response were performed on the combined data, normalized to contralateral and ipsilateral side leading.

ITD increases with eccentricity (Moiseff, 1989): small values of ITD (~0 μs) are associated with frontal directions and larger values of ITD (~200 μs) are associated with the periphery. Thus, we converted ITD to azimuth (Moiseff, 1989) to measure the bias predicted by the mTeg response to ITD.

#### Behavioral data

Behavioral experiments used three different owls, two females and one male. Head orientation and angular velocity were computed offline using image-analysis software (Xcitec, Proanalyst). Individuals analyzing the data remained blind to the location of the sound source until the end. Subsets of the data were analyzed independently by different persons to confirm the analysis was consistent across individuals. Trials where the owls’ bodies moved during the presentation of the stimulus or where the beak disappeared from the field of view of the cameras were discarded. Head movements could be classified as rotation, translation, or fixation, based on angular velocity (Ohayon et al., 2006). Saccades were defined as head rotations with an angular velocity greater than 50 deg.sec^-1^ (Knudsen et al., 1979; du Lac and Knudsen, 1990; Ohayon et al., 2006). When translational movements were included in the analysis the results remained consistent, albeit more variable. We measured saccade latencies as the time from the onset of each presentation of a stimulus to the first video frame where the owl’s head started rotating. Median latency varied across animals (owl L=240 ms; owl F=520 ms; owl P=210 ms). These values are higher than previous reports (Hausmann et al., 2009; du Lac and Knudsen, 1990; Knudsen et al., 1979). The shorter response latency may be attributed to both the motivational and anticipatory state of the owls as most previous studies were conducted with trained owls that were rewarded if the saccade was correct. For each owl, there was no significant difference between the latencies across stimulus conditions (Wilcoxon rank-sum test, p = 0.98). Saccades whose latency was greater than 1 sec were discarded. The criteria for classifying movements as saccades became unreliable when the sound source was close to the front (Knudsen et al., 1979; Hausmann et al., 2009), therefore only head-turns evoked by sources located >10 degrees away from the initial head angle were included in the analysis.

### Modeling

#### Heterogeneous population of neurons in the space-maps

We modeled a space-map population of 500 neurons. To incorporate the overrepresentation of frontal space into the modeled population, the density of preferred direction *X* was drawn from a Laplace prior:

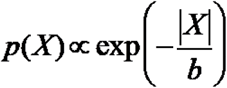

where the scale parameter is *b*=25 (Fischer and Pena, 2011; Knudsen, 1982). First, broadband noise signals between 0.2 and 12 kHz were passed through a gammatone filter-bank with center frequencies ranging from 0.25 kHz to 8 kHz in 0.25 kHz steps. The time constants of the filters were specific to the owl and estimated from Koppl (1997) to model cochlear filters, as described in Fischer et al. (2009). The outputs were cross-correlated and transformed by an exponential input-output function to produce a set of frequency-dependent ITD curves (from 0.5 to 8 kHz). The input to ICx was modeled as a weighted sum of the frequency-dependent ITD curves. The weights were varied with direction to model the direction-dependent frequency tuning in ICx (Cazettes et al., 2014). The weights for each neuron have a Gaussian shape with a center and width selected to match the observed frequency tuning at the preferred direction of the neuron. The weighted sum of the frequency-dependent ITD curves was passed through an exponential nonlinearity to produce the firing rates of Poisson neurons that modeled ICx responses. The responses in OT, which receive point-to-point projections from ICx (Knudsen and Knudsen, 1983), are influenced by feedback inhibition that serves to select the dominant stimulus when multiple peaks of activity are present in the map (Mysore and Knudsen, 2012, 2014). To model OT neurons, we used a simplified version of the model of Mysore and Knudsen (2012, 2014) by multiplying the ICx activity pattern by a rectangular window function to select the dominant peak. The window takes a value of one on an interval that is 28 degrees wide centered at the location of the peak in a smoothed version of the ICx activity pattern. The decoding analyses were applied to the windowed response pattern representing the population activity of the OT.

We hypothesized that a linear transformation of the OT population responses could explain the tunings of neurons in mTeg. Under this hypothesis the mTeg response to ITD should be explained entirely as a linear combination of ITD-tuning curves from OT. Therefore, we modeled the responses in mTeg with a weighted sum of the OT population. To estimate the weights of the convergence, we used discrete Fourier transforms (FFT) of the experimental mTeg ITD tuning curves calculated after offsetting the curves to zero mean, as described previously (Vonderschen and Wagner, 2012). The weights given by the normalized FFT phase spectra of mTeg neurons were applied to OT neurons with equivalent preferred frequency. The modeled mTeg response was therefore given by

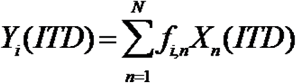

where *X_n_*(*ITD*) is the modeled ITD curve of the n^th^ neuron in OT and *f_i,n_* the weights given by the FFT phase spectra of the i^th^ mTeg neurons at the preferred frequency of the n^th^ OT neuron.

## RESULTS

### ITD tuning in mTeg

To assess the spatial tuning of premotor neurons downstream of the space-maps (i.e, ICx and OT), we conducted single-unit recording of auditory responses in mTeg using metal electrodes. Previous studies have shown that microstimulation of mTeg produces head-orienting responses and eye-movements (Masino and Knudsen, 1993), and that this method could be used to reliably identify this region. Although these studies were successful in establishing mTeg as a region that commands orienting behaviors, sound-evoked responses of these neurons were never explored. To target mTeg we used the same stereotaxic (Masino and Knudsen, 1992, 1993) and microstimulation (Masino and Knudsen, 1990, 1993; Netser et al., 2010) guidelines reported previously. For each recording site, it was verified that: 1) microstimulation at the recording site elicited micro-saccades (Figure 2A,B) and 2) that microstimulation with the same current intensity no longer elicited saccades after the electrode was moved a few hundred microns away from the recording site.

**Figure 2.**
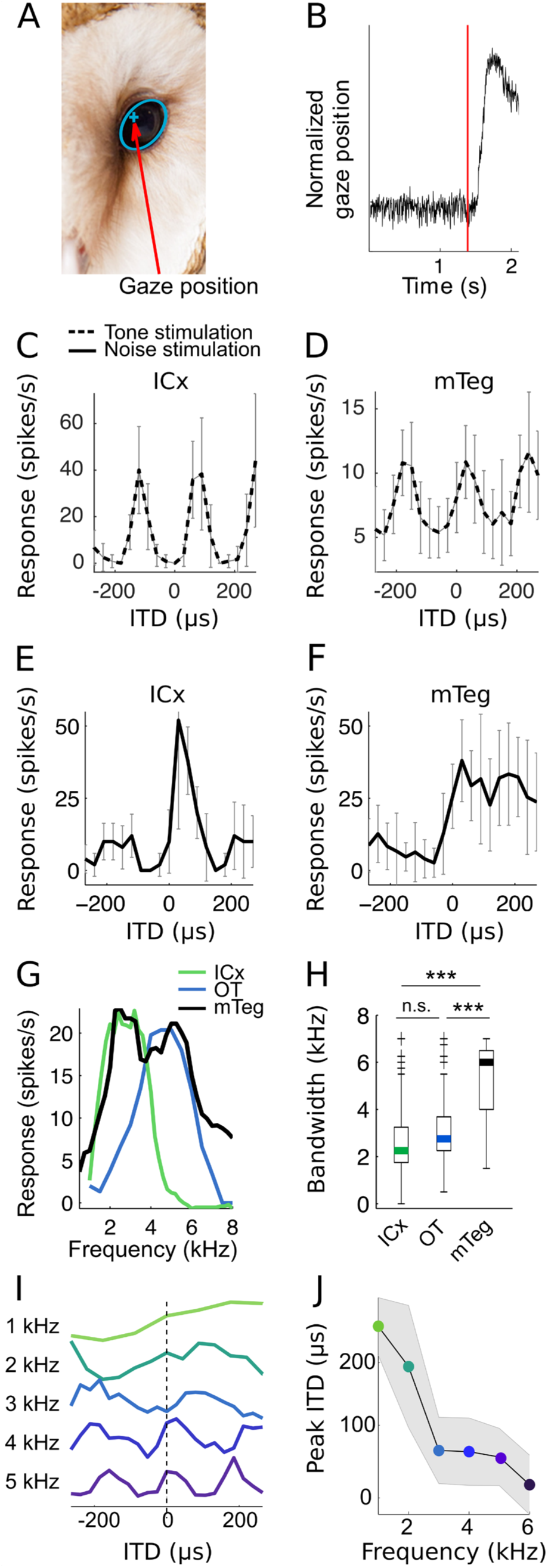
Tuning properties in the space-map and mTeg. (A) An eye tracking system detected the pupil (blue ellipse) and the position of the corneal reflection (blue cross) over time. (B) Microstimulation (red line) through the recording electrode elicited eye movements detected by the eye tracker. (C,D) Example ITD tuning curves (mean and standard deviation across trials) of ICx (C) and mTeg (D) neurons for tonal stimulation. (E,F) Example ITD tuning curves (mean and standard deviation across trials) of ICx (E) and mTeg (F) neurons for broadband noise. (C-F: mean and s.d.). (G) Evidence of frequency convergence from space-map to mTeg. Example frequency tuning curve of an mTeg neuron (black) superimposed with frequency tuning curves of ICx (green) and OT (blue) cells showing the broader frequency tuning in mTeg. (H) Across the samples, mTeg neurons (n=36 from Owls #1, 2 & 3) were significantly more broadly tuned than ICx (n=184 from Owls #1, 2 & 3) and OT (n=39 from Owls #4, 5 & 6) cells (Colored bar represents median, box is 1^st^ and 3^rd^ quartiles, whiskers are 5 and 95%, crosses are greater than 95%, ***p<0.001). (I) Evidence of convergence across ITD channels from space-map to mTeg. Example ITD tuning curves of an mTeg neuron for different frequencies, showing that ITD tuning varied with frequency. (J) The mean and standard deviation of peak ITDs for tonal stimulation across the mTeg sample. The mean peak ITD varied with stimulus frequency in a manner consistent with how ITD and frequency tuning change in the space-map (Figure 1).

Like ICx and OT neurons, mTeg neurons were tuned to ITD. For tonal stimulation, ITD curves in both ICx (Figure 2C) and mTeg (Figure 2D) were periodic, as previously reported in OT (Saberi et al., 1999). However, when broadband noise was used, a notable difference was observed. Unlike neurons in ICx, which display ITD curves for broadband noise with a narrow larger peak flanked by smaller side peaks (Takahashi and Konishi, 1986) (Figure 2E), all mTeg neurons stimulated with broadband noise (n = 36) displayed ITD tuning favoring the contralateral hemifield (one exemple neuron is plotted in Figure 2F). The maxima of these broadband-noise tuning curves were away from the midline on the contralateral side. Although in some neurons there was a slight response decrease at eccentric directions (ITD > 100 μs), in the population average this decrease was not statistically significant from the maximum response (F(6,245) = 0.34, p = 0.92, ANOVA), and thus the general population response in mTeg may be viewed as quasi-sigmoidal.

### Emergence of mTeg tuning

We tested whether the ITD tuning of mTeg neurons was consistent with a convergent projection from OT to mTeg, as proposed for the midbrain network guiding eye saccades (Groh, 2001). If mTeg neurons received convergent input across the space-map from OT, their responses should reflect the varying ITD and frequency selectivity of OT neurons previously observed (Knudsen, 1984; Cazettes et al., 2014) (Figure 1). To test this hypothesis, we investigated two specific predictions for the convergence from OT to mTeg on auditory responses: 1) because frequency tuning varies across the space-map (Cazettes et al., 2014; Knudsen, 1984), a downstream neuron receiving convergent input from OT should have broader frequency tuning; and 2) because preferred ITD co-varies with preferred frequency in ICx and OT (Cazettes et al., 2014; Knudsen, 1984), stimulation with tones will drive activity in only a subset of the cells exciting each mTeg neuron. Thus, convergence from OT to mTeg predicts that mTeg neurons should prefer more frontal ITDs for high frequency sounds and more peripheral ITDs for low frequency sounds.

To test the prediction of broader frequency tuning in mTeg, we measured the half-width of frequency tuning curves in ICx, OT, and mTeg neurons. Frequency tuning curves were significantly broader in mTeg than in ICx and OT neurons (Figure 2G,H, mean ± s.d difference in frequency bandwidth: 2.4 kHz ± 0.3 kHz, Wilcoxon rank-sum test, p < 10^−11^). This result is consistent with a convergence from OT into mTeg.

To test the prediction of covariation of preferred ITD and frequency in mTeg, we measured ITD tuning in mTeg for tonal stimulation. As reported above, the ITD tuning curves of mTeg neurons measured with tonal stimulation were sinusoidal functions with the period determined by the stimulating frequency (Figure 2D,I). Remarkably, unlike in ICx and OT, where neurons prefer the same ITD at all frequencies they respond to, i.e., they display a characteristic delay (Takahashi and Konishi, 1986), mTeg neurons’ preferred ITD varied with frequency (Figure 2I,J). For each cell, the peak ITD shifted towards 0 μs when the stimulus frequency increased (Figure 2J, right, r^2^ = 0.85, intercept=250 μs, slope= −0.042 μs/Hz, p = 0.0019). Moreover, the relationship between preferred ITD and frequency in mTeg neurons matched the relationship between preferred ITD and frequency in the space-map (Cazettes et al., 2014) (r^2^ = 0.75, intercept=284 μs, slope= −0.045 μs/Hz, p<10^−55^), where frontal neurons, tuned to ITDs near zero, prefer higher frequencies than peripheral neurons tuned to large ITDs (Figure 1) (Cazettes et al., 2014; Knudsen, 1984). These results provide physiological evidence consistent with a convergence from OT into mTeg.

### Weighted convergence from space-map to mTeg

It was shown above that mTeg tuning to ITD and frequency are consistent with a convergence that takes place between the space-map in OT and mTeg. We next asked whether the mTeg responses can be explained by a weighted linear convergence from space-map neurons, a processing scheme that has been proposed for the emergence of motor commands (Groh, 2001; Salinas, 2006). For this, a bank of ITD tuning curves of a population of OT neurons was modeled following criteria from published tuning properties of space-map neurons. Specifically, cells tuned to frontal ITDs prefer higher frequencies and therefore display narrower ITD tuning (Cazettes et al., 2014; Knudsen, 1984) (Figure 3A). This modeled OT population was the same used previously for Bayesian decoding analyses (Cazettes et al., 2016). Modeled OT tuning curves were then weighted (Figure 3B,C) and summed (Figure 3D) to simulate the weighted convergence. To estimate the weights at each frequency/ITD channel converging onto mTeg cells, we used linear decomposition by Fourier analysis of ITD tuning curves for broadband noise of mTeg neurons. It has been shown that this method can be used to efficiently assess the neurons’ frequency responsiveness (Vonderschen and Wagner, 2012) (Figure 3B). Here, a higher power at a specific frequency in the Fourier decomposition of the broadband-noise ITD response of an mTeg neuron corresponds to a higher weight applied to the projecting OT neurons corresponding to that frequency (Figure 3A-C). The weighted OT ITD-tuning curves were then summed and the output was compared to the actual ITD tuning of mTeg neurons on a cell-by-cell basis (Figure 3D). We observed strong correlations between modeled and measured mTeg ITD-tuning curves (Figure 3E, pairwise linear correlation, p<0.05 for 34 out of 36 neurons, median variance explained = 70%), indicating that ITD tuning of mTeg neurons to broadband sound could be predicted from the weighted convergence of the space-map population. These results indicate that mTeg responses are consistent with a weighted linear convergence from the midbrain space map, a plausible mechanism in neural processing (Groh, 2001; Salinas, 2006).

**Figure 3.**
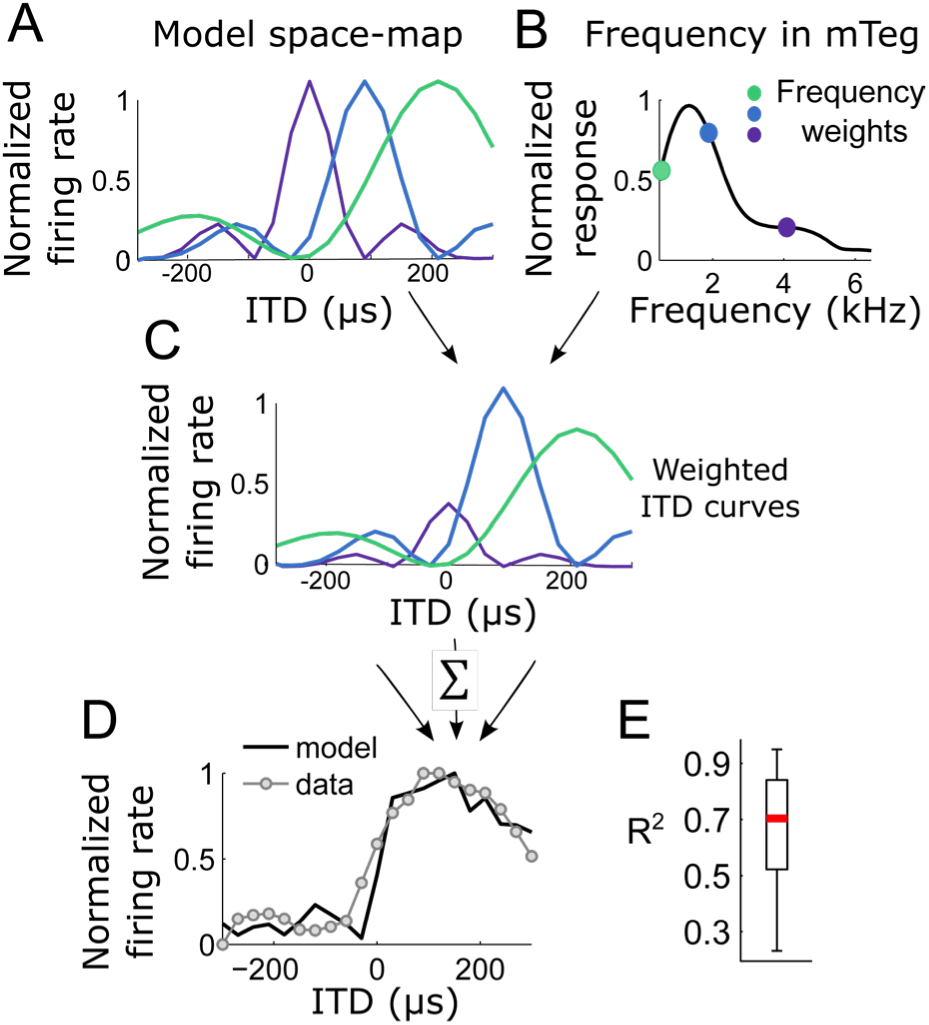
Weighted convergence from the space-map into mTeg. (A) Model of frequency-dependent ITD tuning curves in the space map. Neurons tuned to small ITDs preferred high frequencies (purple), neurons tuned to intermediate ITDs preferred intermediate frequencies (blue) and neurons tuned to larger ITDs preferred low frequencies (green). (B) The weights for each frequency (normalized from 0 to 1) were estimated from the Fourier decomposition of responses of mTeg neurons to broadband sound, on a neuron-by-neuron basis. (C) Space-map ITD tuning curves (A) scaled by the weights estimated in (B) and summed to simulate the convergence. (D) Example experimental ITD-tuning curve in mTeg (gray dotted line) and the output of the convergence model with the respective weights. (E) The boxplot shows the median and quartiles of the coefficient of determination (r^2^) between modeled and measured mTeg ITD curves across the sample.

### Emergence of an adaptive orientation estimate in mTeg

We next examined whether mTeg responses predicted behavior when the reliability of the ITD cue changed. To manipulate the reliability of the ITD cue, the similarity of the sound at the left and right ears was varied by adding uncorrelated noise to reduce binaural correlation (BC). Lowering BC increases the sensory noise. Previously reported data shows that the owl’s localization is increasingly biased toward zero degrees when BC decreases (Figure 4A, reproduced from Saberi et al. (1998)). To compare the mTeg responses with the owl’s localization data, we plotted the mean mTeg responses as a function of BC for the same four ITDs used in the published behavioral experiments. These mTeg responses were well correlated with behavior (Figure 4B, mean r^2^ for the four stimulus directions = 0.84, p=0.017).

**Figure 4.**
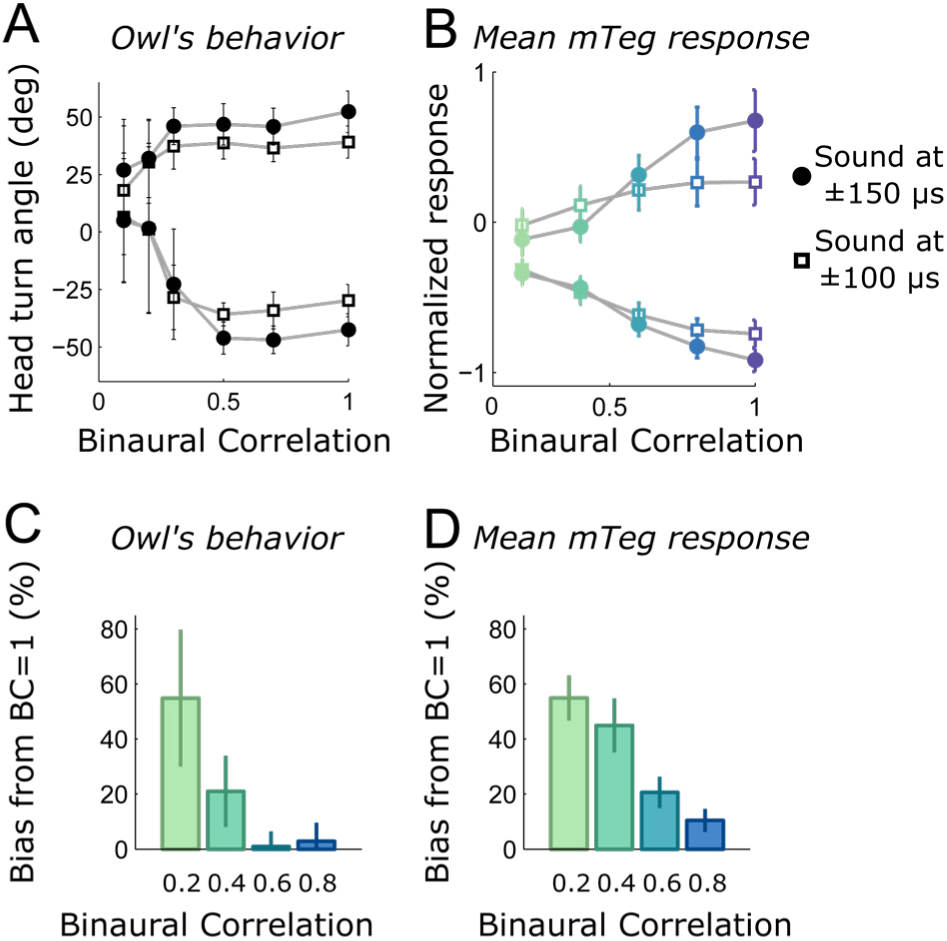
The owl’s behavioral bias under varying levels of noise is consistent with mTeg responses. (A) Mean and standard deviation of the owl’s head orienting response as a function of BC (modified from Saberi et al. 1998). Behavioral responses to four different ITDs are shown (actual values indicated on the right). The centrality bias increases as BC decreases. (B) Mean mTeg response across BC for similar ITDs as in (A). (C,D) Behavioral (C) and average mTeg response changes as a function of BC, plotted as percent of the BC = 1 condition. To quantify the bias toward the center in both behavior and neural responses, the mean difference between responses at BC=1 and other BCs were computed at the same four directions shown in (A) and (B). Error bars represent s.e.m.

Finally, we examined whether the bias in mTeg responses and the owl’s behavioral bias matched as BC changed. To compare the bias in mTeg responses and behavior we computed the mean difference between responses at BC= 1 and at lower BCs (Figure 4C, D). The average mTeg response predicted an increased bias at lower BCs (Figure 4D) that displayed a similar trend as the orienting behavior of the owl (Figure 4C).

Overall, the correlation between mTeg neural responses and behavior across BCs supports the hypothesis that mean mTeg responses approximate an estimate of sound direction that changes adaptively when the sensory input is degraded.

### Testing the network architecture by manipulating the owl’s behavioral bias

To further test the weighted convergence model, we aimed to selectively manipulate the activity in different parts of the space-map and verify whether these manipulations yielded consistent changes in mTeg responses and the owl’s behavior. Our hypotheses were based on previously published evidence showing that frontal and peripheral space-map neurons are tuned to high and low frequencies, respectively (Cazettes et al., 2014; Knudsen, 1984). Therefore, stimulation with a sound containing only high frequencies should enhance the response of frontal space-map neurons relative to peripheral neurons, with the converse occuring for stimulation with low frequencies (Figure 5A). Hence, sound localizing behavior should display stronger bias for the front for high-frequency than for low-frequency sounds. Moreover, if behavior is driven by the mTeg responses, then the difference in mTeg responses to high and low frequency sounds should match differences in behavioral localization bias for high and low frequency sounds.

**Figure 5.**
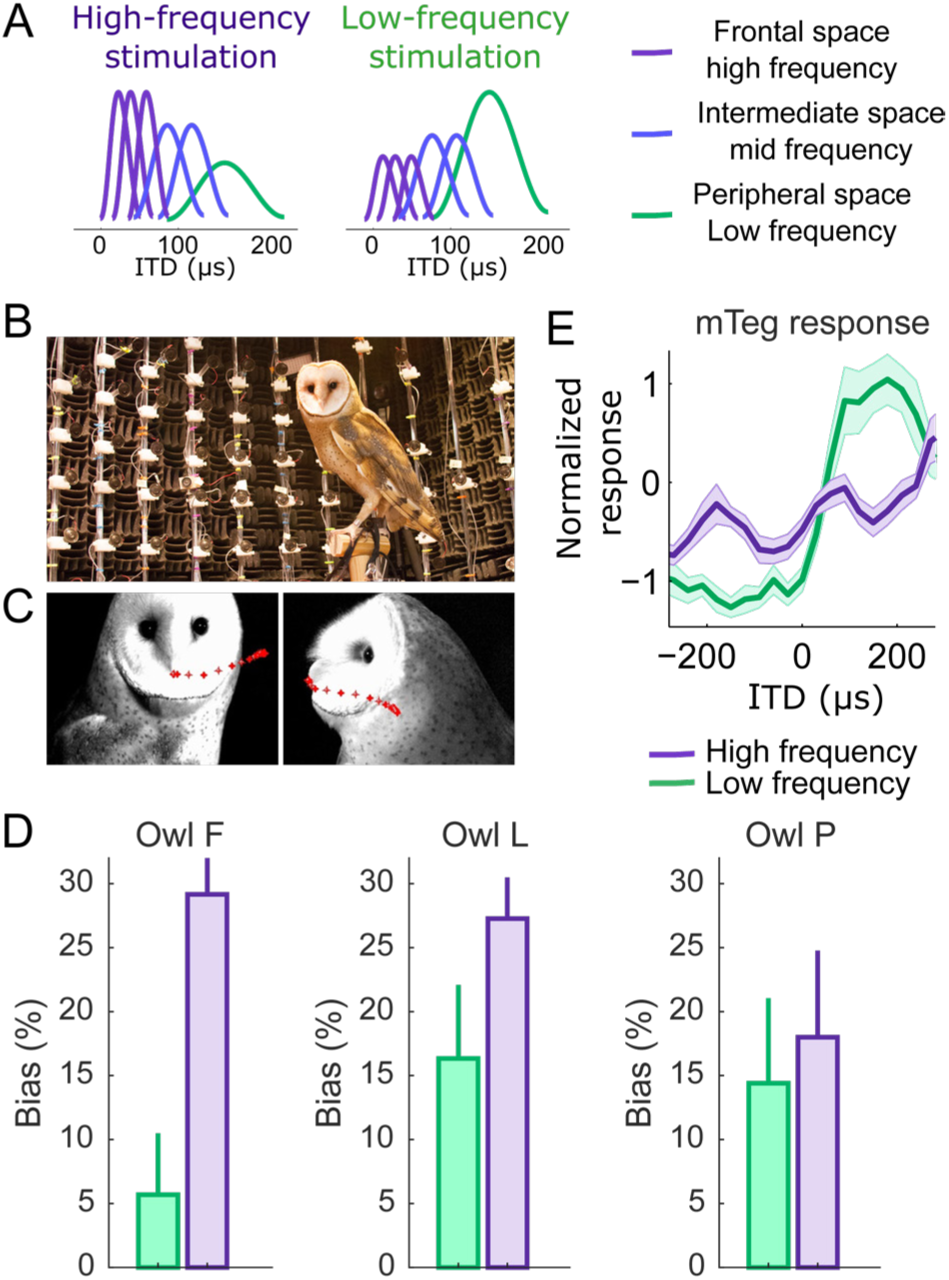
Changes in the owl’s behavioral bias and mTeg responses induced by manipulating stimulus frequency. (A) Schematic population of space-map neurons whose relative response rate varies with the frequency of the stimulating sound. Stimulation with high frequencies (middle) excites more strongly the frontal neurons, which prefer higher frequencies. Conversely, stimulation with low frequencies (bottom) excites more strongly the peripheral neurons. (B) Owls were trained to stand on a perch at the center of a speaker array. (C) Images captured by two high-speed infrared video cameras at different angles. The owl’s head was tracked as it moved in space. Red dotted lines indicate the trajectory preceding the shown images. (D) Mean and s.e.m of the bias of head-orienting responses from three barn owls (F, L and P) presented with high-frequency and low-frequency bandpass noise. (E) Mean ITD tuning across mTeg neurons for low-frequency (green) and high-frequency (purple) bandpass noise.

To test whether the owl’s behavioral bias was indeed consistent with these predictions, we measured non-reinforced head-turns triggered by high and low-frequency band-pass as well as broadband sounds in behaving barn owls. Training was limited to standing on a perch at the center of a speaker array; in this way we avoided potential influences of over-training (Hazan et al., 2015) (Figure 5B). The owl’s natural tendency to orient the head toward sounds was tracked with high-speed infrared cameras (Figure 5C). Head-saccades were measured from any initial head angle to avoid positional biases. To control for the confound between centrality bias and unwillingness of the barn owls to turn the head for sounds close to the center of gaze, sound sources within 10 degrees or less from the center of gaze at the time of sound presentation were not included in the analysis. We then estimated the average bias as the mean difference from the actual sound source direction. Using this paradigm, owls underestimated peripheral directions for broadband noise (the average bias over all eccentric directions was 21 deg), matching reports in previous studies using owls trained to fixate at the front and motivated to respond through reward (Hausmann et al., 2009; Knudsen et al., 1979). Remarkably, owls showed significantly stronger underestimation when localizing high frequency sounds than low frequency sounds (p =0.00004, permutation test; average bias at high frequency: 27 deg; average bias at low frequency: 13 deg, see Figure 5D for the performance of each owl). Thus, the amount of behavioral bias could be adjusted upward and downward by targeted manipulations of the sensory input.

We then measured mTeg responses to the same high and low frequency sounds used in the behavioral experiments. We found that mTeg responses were consistent with the behavior, displaying significantly lower firing rate at peripheral ITDs for high than low frequencies (Figure 5E, F(2,54) = 9.35, p = 0.0003, ANCOVA). Specifically, the population mean mTeg response to ITD with high frequency sound was 30% lower than mTeg response to ITD with low frequency sound. In summary, the consistency between the mTeg neural responses and behavioral bias observed during stimulus’ frequency manipulations indicates, independently from the BC manipulations, that mTeg responses could account for the behavioral bias by integrating across the space-map population.

## DISCUSSION

This work provides evidence of how the midbrain map of auditory space may be read out by downstream neurons to synthesize a command of orienting behavior that captures sensory reliability. It has previously been shown that the selectivity of space-map neurons reflects cue reliability (Cazettes et al., 2014) and was proposed that this property may be a crucial parameter for guiding an adaptive behavioral bias (Cazettes et al., 2016). However, how exactly changes in space-map neurons’ selectivity are translated into an adaptive behavior was left unanswered. The present study shows how the space-map population may be read-out by downstream neurons to guide behavior in situations of uncertainty. Here, we demonstrated that this processing is consistent with a weighted convergence from the OT to mTeg, generating a conversion from a map representation to a population rate-code that can drive orienting behavior in a manner that reflects sensory reliability. Based on these mechanistic findings, predictions were made about how the adaptive behavioral bias would change by controlling stimulus frequency. These predictions were confirmed with behavioral experiments, allowing further demonstration of the network architecture by manipulations at the encoding level. Therefore, we show here how sensory statistics can be biologically decoded into an adaptive signal controlling head saccade.

### Sensory-motor integration

Critical findings of this study are the evidence of convergence from OT onto mTeg and the effect this convergence has on the ITD tuning in mTeg. Convergence from the space-map is supported by the broader frequency tuning of mTeg neurons and their frequency-dependent ITD tuning, which matches the ITD-dependent frequency tuning that was previously reported in ICx and OT (Cazettes et al., 2014; Knudsen, 1984). The remarkable consistency of how best ITD changes with frequency in mTeg with how best frequency changes with ITD in ICx (Cazettes et al., 2014) and OT (Knudsen, 1984) indicates that convergent projections of neurons from the space-map can explain the tuning in mTeg. As a consequence of this convergence, mTeg neurons lacked the classic characteristic-delay described for neurons of the inferior colliculus (Takahashi and Konishi, 1986). This is also consistent with the different ITD tuning of ICx and OT neurons, compared to mTeg cells. Whereas ICx and OT responses display peaked and largely symmetric rate-ITD curves around a prominent preferred ITD for broadband sound, mTeg responses are largely asymmetric with maximal responses uniformly away from zero ITD.

A fundamental question in sensorimotor integration is the neural code controlling movement (Todorov, 2000; Salinas, 2006; Overduin et al., 2012; Joshua and Lisberger, 2014). Previous studies have reported correlations between the firing rate of sensorimotor neurons and the kinematics of movements (Todorov, 2000). Here we propose that the premotor neurons in mTeg encode direction in the population average firing rate, which, through weighted convergence, captures the statistics of the sound localization cues that are represented in the upstream space-map. This is consistent with the idea that mTeg neurons control muscle synergy rather than single muscles independently (Masino and Knudsen, 1990; Overduin et al., 2012; Joshua and Lisberger, 2014), which has been proposed for explaining why correlations between neuronal responses and movement properties are commonly observed (Todorov, 2000).

The dorsal and medial tegmental areas are composed of multiple nuclei that receive direct projections from the OT. Microstimulating these areas yields head-saccades in different directions (Masino and Knudsen, 1993). Therefore, populations of mTeg neurons in distinct nuclei are likely to have different tuning properties. Here, we focused on the horizontal coordinate of sound direction, which in barn owls is specifically linked to the coding of ITD (Moiseff, 1989). Hence, we targeted mTeg neurons with clear tuning to ITD throughout mTeg. Because of the limited number of owls and the need to maximize the data collection, euthanasia of birds used for recordings was not conducted, and we relied on physiological methods and microstimulation to identify ITD-tuned mTeg cells. While the purpose of this study was to elucidate the network underlying the emergence of an adaptive motor command for head lateralization, we did not attempt to completely explain the anatomical pathway supporting head-orienting responses. Further experiments are needed to explore the heterogeneous and likely complex properties of mTeg neurons that participate in guiding eye and head movements in different directions and contexts.

### Implementation of an adaptive behavior

In general, behavioral errors are composed of systematic errors (biases) and precision errors (variance). Previous studies have focused on behavioral biases to understand the underlying neural code (Schwartz et al., 2007; Jazayeri and Shadlen, 2010; Summerfield and Lange, 2014; Wei and Stocker, 2012, 2015). Focusing on the bias itself can provide information about the strategy the brain uses to produce adaptive behavior. Although both bias and variance contribute to the magnitude of errors, they provide different insights into how errors may occur in judgments. While the variance may reflect the level of confidence in either the evidence or the estimate, the systematic nature of the bias and its direction can be interpreted as the manifestation of a prior (Fischer and Pena, 2011; Girshick et al., 2011; Cazettes et al., 2016; Wei and Stocker, 2015) or a cost function (Kording and Wolpert, 2004) that constrains the judgment. Understanding the neural response properties that emerge from the underlying circuitry in the owl’s sound localization pathway made it possible to manipulate the behavioral bias using specific sensory stimuli. Thus a simple and biologically plausible mechanism of weighted convergence can explain how the neural code for sound localization can be translated into an adaptive behavioral command. These findings may also be testable in other species where auditory biases have been observed, including humans (Zahn et al., 1978; Populin and Yin, 1998; Nodal et al., 2008).

## Conflict of interest

The authors declare no competing financial interests.

## Acknowledgements

This work was funded by NIH grants DC012949 and DC007690. We thank Kourosh Saberi for providing the data used in Fig. 4A and Adam Kohn for comments on the manuscript. Clifford Keller helped hand-raise the owlets and Daniel Munch participated in the blind analysis of behavior.

